# Estimating universal mammalian lifespan via age-associated epigenetic entropy

**DOI:** 10.1101/2024.09.06.611669

**Authors:** Juan José Alba-Linares, Juan Ramón Tejedor, Agustín F. Fernández, Raúl F. Pérez, Mario F. Fraga

**Affiliations:** Cancer Epigenetics and Nanomedicine Laboratory, Centro de Investigación en Nanomateriales y Nanotecnología-Consejo Superior de Investigaciones Científicas (CINN-CSIC), Universidad de Oviedo, 33011 Oviedo, Spain; Instituto de Investigación Sanitaria del Principado de Asturias (ISPA), 33011 Oviedo, Spain; Instituto Universitario de Oncología del Principado de Asturias (IUOPA), Universidad de Oviedo, 33006 Oviedo, Spain; Centro de Investigación Biomédica en Red de Enfermedades Raras (CIBERER), Instituto de Salud Carlos III (ISCIII), 28029 Madrid, Spain; Departamento de Bioquímica y Biología Molecular, Facultad de Veterinaria, Universidad Complutense de Madrid, 28040 Madrid, Spain; Departamento de Biología de Organismos y Sistemas, Área de Fisiología Vegetal, Universidad de Oviedo, 33071 Oviedo, Spain

**Author notes:** Correspondence to: Raúl F. Pérez, Mario F. Fraga.

**Keywords:** lifespan, DNA methylation, aging, mammalian, epigenetic entropy

## Abstract

Loss of epigenetic information has been proposed as a potential driver of mammalian aging. However, its contribution to the well-documented variation in lifespan estimates among mammals remains to be elucidated. In this study, we examined DNA methylation entropy patterns at evolutionarily conserved CpG sites across multiple mammalian species to quantify age-associated epigenetic information loss. We found that longer-lived species tend to accumulate fewer CpGs exhibiting increased methylation noise over time, irrespective of whether these changes arise from hyper- or hypomethylation mechanisms. Importantly, the rate of epigenetic entropy gain declines in a linear fashion with species’ maximum lifespan, pointing to the existence of a universal constraint on mammalian longevity, estimated to lie in the vicinity of 220 years. We further demonstrated that this relationship and its associated limit were independent of species and sample selection, as well as phylogenetic relatedness, and remained robust across different scenarios of lifespan estimation uncertainty. Collectively, this work highlights the maintenance of epigenetic information as a key factor in explaining lifespan differences among species and proposes a universal maximum limit to natural mammalian longevity.

## Introduction

The species that comprise the tree of life encompass a vast array of organisms with unique life traits covering a dynamic range of biological features such as body mass, metabolic rate, and developmental and reproductive patterns. Nevertheless, although maximum lifespan is species-specific(1), the molecular mechanisms underlying interspecies differences remain unidentified.

In this regard, many theories have been proposed to elucidate the evolutionary origins of aging and lifespan control(2). At the molecular level, stochastic (enzymatic errors, reactive by-products) and deterministic (genetic programs, environmental conditions) factors have been proposed to drive an accumulation of molecular damage and disorder ultimately leading to aging(3, 4). For instance, loss of cellular identity is a hallmark of aged tissues, and appears to be driven by the convergent transcriptional activation of tissue-specific genes due to heterochromatin erosion with age(5–7). Recently, loss of epigenetic information has been put forward as a holistic framework to explain possible mechanistic processes underlying aging(8). In this scenario, genome-wide epigenetic erosion due to the repair of DNA damage is sufficient to trigger aging *in vivo*, including cellular dedifferentiation and acceleration of DNA methylation clocks, which can be reversed by cellular reprogramming using Yamanaka factors(9). Nevertheless, whether the accumulation of biological noise may explain lifespan differences across species remains unresolved.

In this context, Cagan and colleagues revealed that loss of genetic information, in terms of somatic mutation rate per year, scales with lifespan in mammals(10), and DNA methylation seems to follow a similar law of exponential decay(11), primarily at PRC2-bound bivalent promoters(12), which also tend to gain methylation with age(13). In the same vein, PRC2 domains experience intense concurrent entropy gain with age, which can be reversed through long-term partial reprogramming(14). Additionally, regional methylation disorder in PRC2-target genes shared across four mammals exhibited a deceleration with longer lifespan(15). However, the role of PRC2-independent hypomethylation mechanisms still requires further investigation.

In DNA methylation array studies, epigenetic noise has been typically measured using various metrics, with Shannon entropy (see Materials and Methods) being among the most common(16). Notably, epigenetic entropy has been shown to be as important as average methylation in understanding core biological processes such as development(17). Consequently, we hypothesized that epigenetic entropy gain over the course of a lifetime is a species-specific characteristic, and that a lower accumulation of noise corresponds to a longer maximum lifespan. Therefore, our work aimed to address fundamental questions regarding the relationship between DNA methylation and mammalian maximum lifespan limits: 1) Can both DNA methylation gain and loss contribute to age-related increases in entropy that scale with species’ maximum lifespan? 2) If so, can entropy patterns be used to estimate the upper limit of maximum lifespan achievable by any mammalian species?

To address the aforementioned questions, we conducted a comprehensive investigation into epigenetic entropy gain dynamics in CpG sites from the Mammalian array(18–20) across 18 species that met the necessary criteria for rigorous statistical analysis (see Materials and Methods). The recent discovery of tissue-to-tissue disparities in the prediction of species’ lifespan via novel epigenetic clocks has led to the identification of blood as the optimal tissue for consistently yielding the highest lifespan estimations across species(21). Accordingly, our approach entailed the examination of blood samples from diverse taxonomic orders that encompassed the majority of maximum lifespan observations documented in mammals.

## Results

### Long-lived mammalian species show less DNA methylation noise accumulation with age

To investigate whether epigenetic entropy gain with age correlates with maximum lifespan, we first focused on selecting comparable mammals with sufficient lifetime blood DNA methylation profiles (see Materials and Methods). 18 species (3,107 annotated blood samples) satisfied our selection criteria while ensuring a balanced and adequate representation of distinct phylogenetic orders (Fig. 1A), with 4 primates, 4 rodents, 4 carnivores, 4 artiodactyls and 2 elephantids spanning a wide lifespan range both within and between clades.

**Figure 1.**
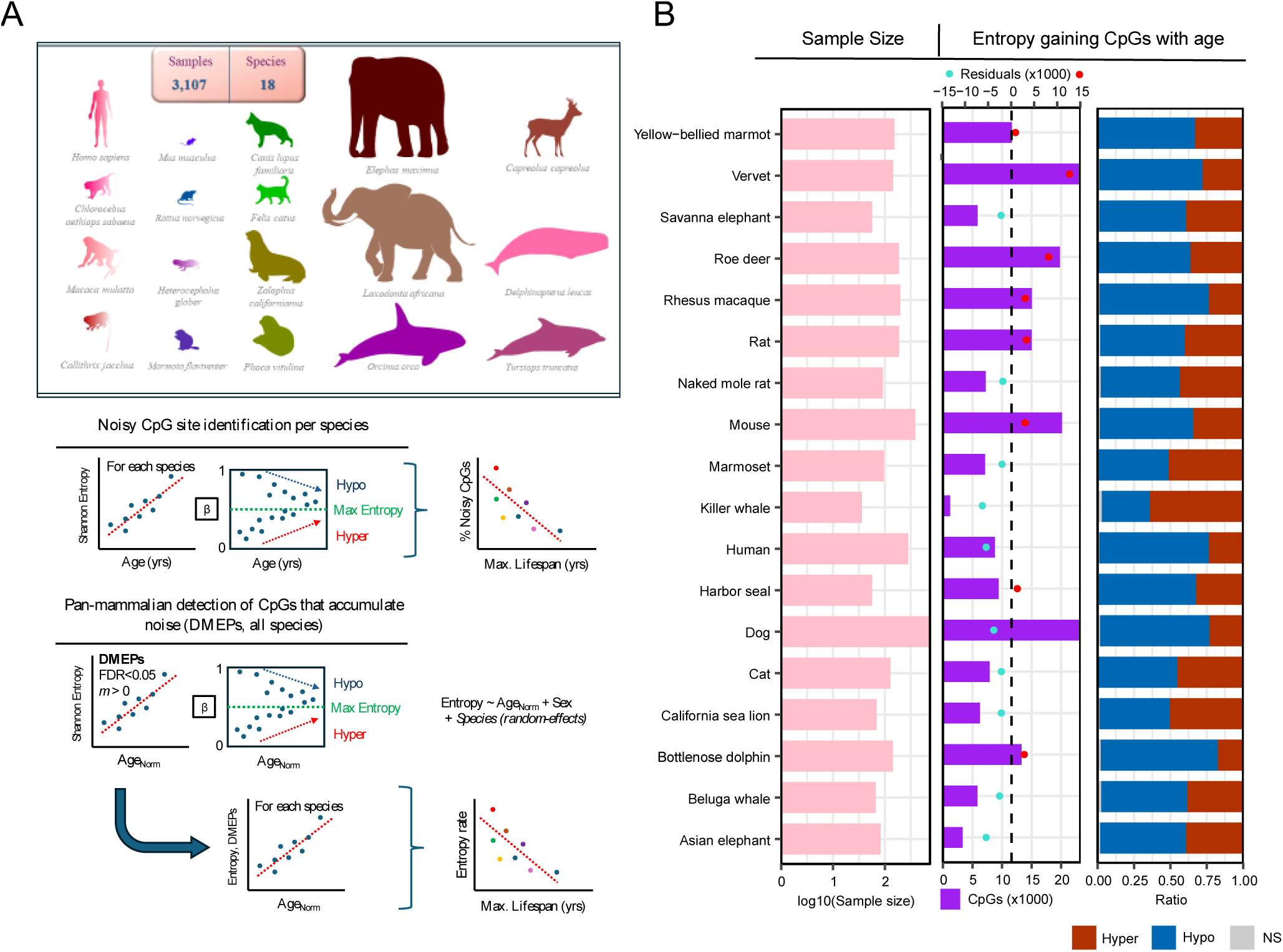
Entropy accumulation with age results from bidirectional loss-gain methylation mechanisms. (A) Overview of the study design and computational workflow. First, CpG sites accumulating age-associated epigenetic noise were identified in each species to assess whether their absolute numbers correlated with species’ maximum lifespan. Next, a pan-mammalian model was built using all available data and normalized ages to detect pan-species differentially methylated entropic positions (DMEPs). Finally, the rate of entropy gain at these DMEPs was correlated with species’ maximum lifespan. (B) Barplots depicting: left, the sample size distribution per species; middle, the number of CpGs that become noisier with age per species, along with the residuals obtained after adjusting for sample size effects; right: the relative distribution of these CpG sites based on the direction of methylation change.

Next, we used linear regression models to examine how the dynamics of epigenetic noise behave during aging at the single-CpG level for each species (Fig. 1B). The number of CpG sites that accumulate disorder with age (FDR<0.05, logFC>0) varied considerably across mammals, but primarily resulted from hypomethylation mechanisms in the majority of species, which is expected given that most CpG sites displayed high average methylation levels in the Mammalian array.

In view of this interspecies variability, we observed that the number of significant hits was inversely proportional to the maximum lifespan of each mammalian species (Fig. 2A: *R =* −0.5530, *P =* 0.0173). Furthermore, this association was maintained when segregating entropy-gaining CpGs based on the direction of their average DNA methylation change (Fig. 2B, hyper: *R =* −0.7840, *P =* 0.0001; Fig. 2C, hypo: *R =* −0.4299*, P =* 0.0750). To circumvent potential biases associated with unequal numbers of samples per species, which can modify the statistical power of detecting significant hits, we adjusted the number of entropy-gaining CpGs per species by regressing this variable against corresponding sample sizes and extracting the residuals. This approach confirmed the observed negative trends while adjusting for this potential confounding factor (Supplementary Fig. 1A-C, see Materials and Methods). It is our contention that the strength of the anticorrelation may be greater for DNA hypermethylation than for hypomethylation because the former involves fewer CpGs in the majority of species (Fig. 1B).

**Figure 2.**
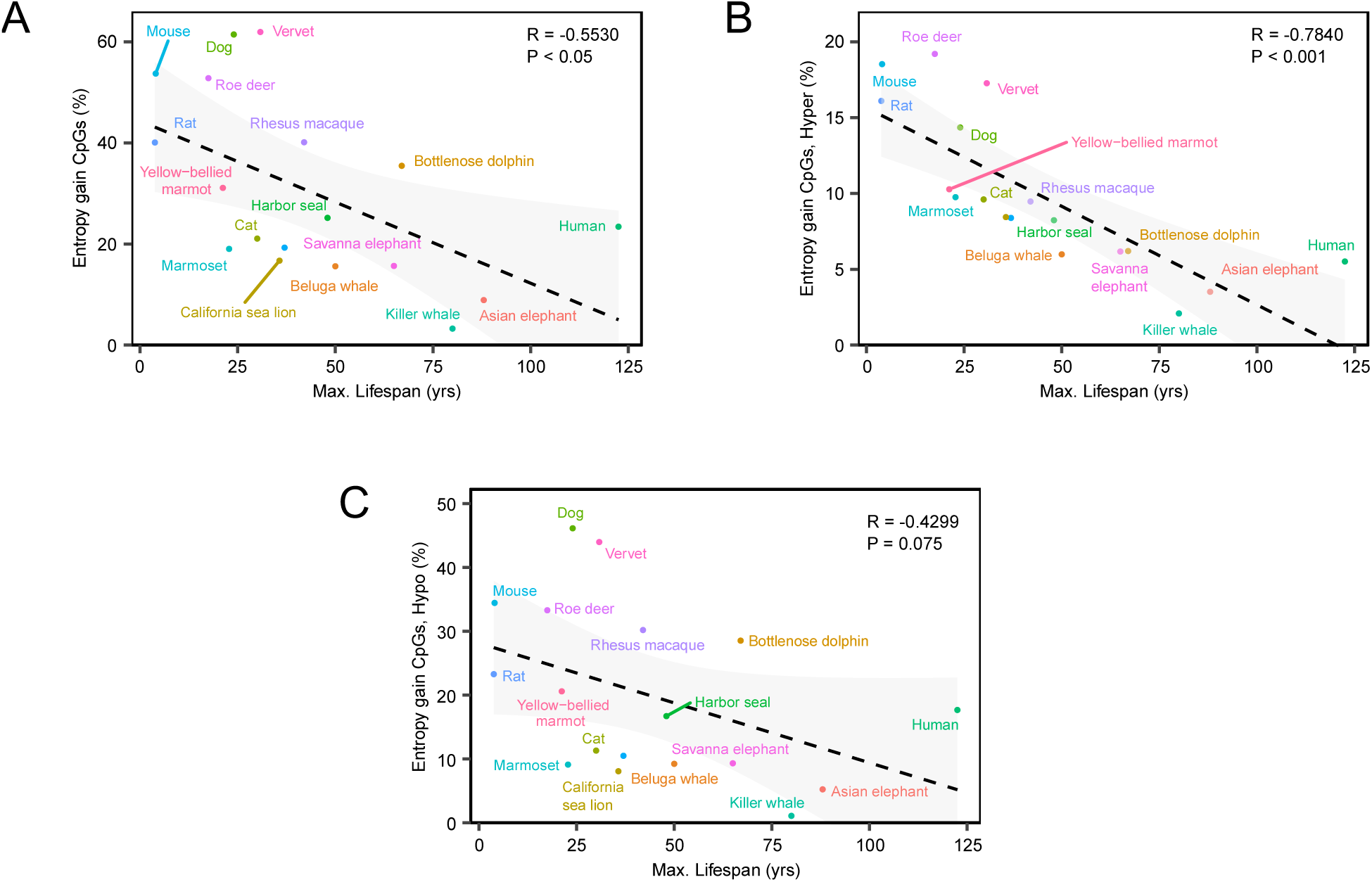
The number of CpGs that acquire epigenetic noise during aging declines with species’ maximum lifespan. **(A-C)** Scatter plots showing the percentage of CpGs gaining epigenetic entropy with age **(A)**, further separated into CpGs undergoing hyper- **(B)** and hypomethylation **(C)**.

To further validate this discovery, we replicated our analyses after removing probes with non-bimodal distributions across the 18 species, as these could represent conflicting or failed signals (Supplementary Fig. 2, see Materials and Methods). With this stricter selection of 18,268 probes (out of 37,488, 48.73%), we observed even stronger anticorrelations between maximum lifespan and the number of bimodal, entropy-gaining CpG sites per species (Supplementary Fig. 3A, all: *R =* − 0.5769, *P =* 0.0122; Supplementary Fig. 3B, hyper: *R =* − 0.7935, *P =* 8.51 × 10^−5^; Supplementary Fig. 3C, hypo: *R =* − 0.4570, *P =* 0.0566). The results presented herein demonstrate that the CpG sites that gain epigenetic entropy with age, irrespective of the direction of methylation change, contain information capable of tracing maximum lifespan trajectories in mammals.

### Species-specific entropy accumulation rates reflects maximum mammalian lifespan

After observing the association between maximum lifespan and the numbers of CpGs gaining noise with age in each species, we sought to explore pan-species patterns of entropy gain during aging in the Mammalian array using a unified model across all species. This strategy allowed us to test whether the rate of noise accumulation in each species was linked to their respective lifespan, while accounting for potential biases such as uncertainty in species-specific entropy rate estimates, possible maximum lifespan misestimations, phylogenetic relationships, and differences in normalized age distributions.

To do so, we first built a multivariate linear mixed model, incorporating species and sex as covariates, and define DMEPs as CpG sites that gain epigenetic entropy as a function of normalized age (FDR<0.05, logFC>0, see Materials and Methods), an approach that enabled the simultaneous analysis of the 18 selected species in our generalized model (Fig. 1A). We observed that the overwhelming majority of CpG sites (21,876) were defined as DMEPs (Supplementary Table 1), far exceeding the sum of pan-species CpG sites that lost epigenetic entropy (7,459) or remained stable (8,153) (Fig. 3A). Notably, a higher proportion of these DMEPs (68.6%) originated from DNA hypomethylation processes.

**Figure 3.**
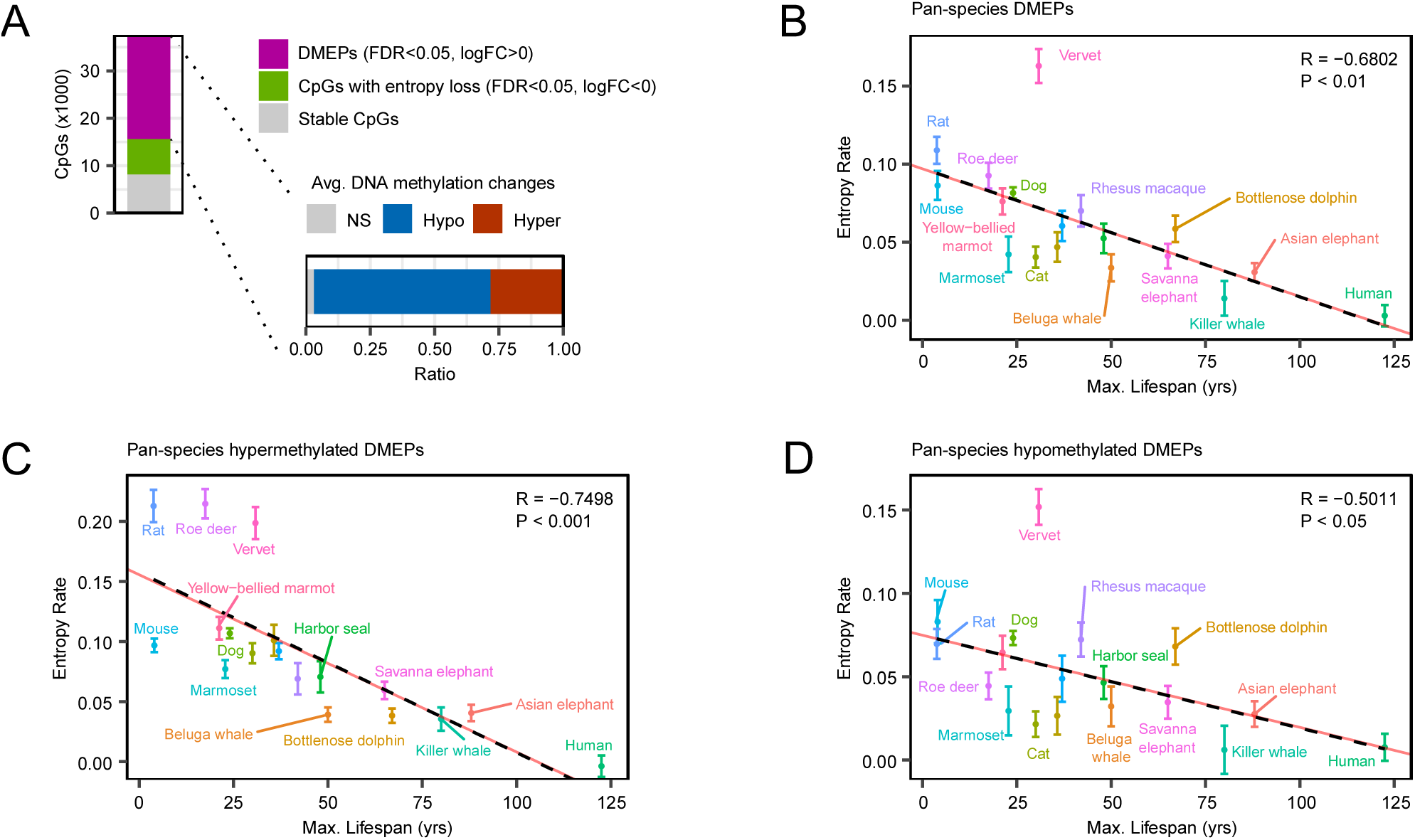
The rate of methylation entropy gain throughout lifetime tracks species’ maximum lifespan. **(A)** Barplot classifying mammalian array CpGs according to their contribution to entropy gain or loss across species. DMEPs are further grouped according to the direction of average methylation change. **(B-D)** Scatter plots showing the linear decrease (dashed black line) in epigenetic entropy gain rate with age as a function of species’ maximum lifespan **(B)**, distinguishing DMEPs undergoing hyper-**(C)** and hypomethylation **(D)** processes. Standard errors for each rate are included, along with meta-analysis regression lines (shown in red).

We then examined the rate of epigenetic entropy gain with normalized age at these DMEPs for each species and its relationship with each species’ maximum lifespan. Importantly, we observed a robust, inversely proportional linear relationship with maximum lifespan in mammals (Fig. 3B: *R =* −0.6802, *P* = 0.0019). Again, this negative correlation was also evident when examining DMEPs derived exclusively from DNA hypermethylation (Fig. 3C: *R =* −0.7498, *P* = 0.0003), or from hypomethylation mechanisms (Fig. 3D: *R =* −0.5011, *P* = 0.0341). A meta-regression model accounting for each species’ entropy rate uncertainty further validated these observations with stronger significance (Fig. 3B-D: *P_meta_* = 0.0001, 4.87 × 10^−6^ and 0.0195 for all, hypermethylation and hypomethylation DMEPs, respectively).

Accurately estimating the maximum lifespan of a species is challenging, as many ecological factors and sampling biases can affect the recording of long-lived individuals in their native settings(22). To address this potential issue, we replicated our model 10,000 times using resampling at both the individual and species level, while simultaneously introducing a gradient of fixed relative errors in maximum lifespan values (±1% to ±50%, see Material and Methods). With this strategy, we observed that introducing even a ±15% misestimation error still maintained a significant association between entropy rates and maximum lifespan in >85% of the bootstrap replicates (Supplementary Fig. 4A). Furthermore, considering a more realistic scenario of systematic maximum lifespan underestimation (Materials and Methods) led to the preservation of associations in >85% of bootstrap replicates with up to +60% positive additive error in lifespan (Supplementary Fig. 4B). This latter finding is of particular importance because our study focuses on studying the maximum lifespan a species is biologically capable of achieving – one that is at least equal to, and possibly greater than, the maximum lifespan recorded for any individual.

In addition to accounting for potential maximum lifespan misestimations, we also tested our model using time-to-sexual maturity, which may be a more reliably characterized demographic trait for longevity studies(23, 24). Subsequently, we found significant and very similar associations between entropy rates and species’ time-to-sexual maturity for all DMEPs (Fig. 4A-C, DMEPs: *R = −*0.6087, *P =* 0.0073; hyper-DMEPs: *R= −*0.7065, *P =* 0.0010; hypo-DMEPs: *R = −*0.4241, *P =* 0.0794), even more significant when accounting for entropy rates uncertainties (DMEPs: *P_meta_=*0.0015; hyper-DMEPs: *P_meta_=*5.10 × 10^−5^; hypo-DMEPs: *P_meta_=*0.0602).

**Figure 4.**
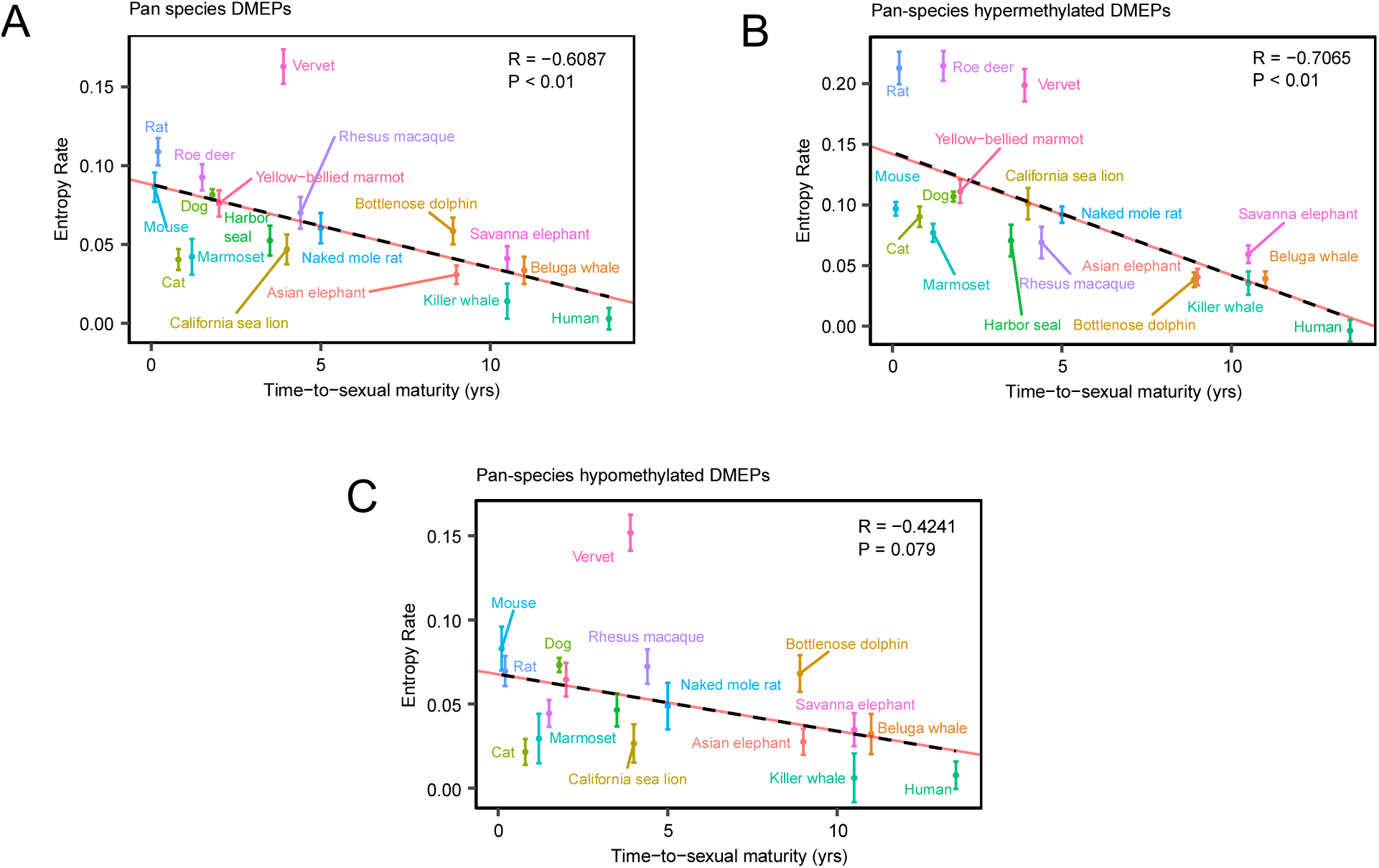
Species with delayed sexual maturity show lower rates of epigenetic entropy gain with age. (A-C) Scatter plots showing the linear decrease in epigenetic entropy gain rate with age as a function of species’ time-to-sexual maturity (A), distinguishing DMEPs undergoing hyper- (B) and hypomethylation (C) processes. Standard errors for each rate are included, along with meta-analysis regression lines (shown in red).

Although unlikely, we also explored the possibility that the observed association between entropy rates and maximum lifespan could be influenced by phylogenetic relationships between the analyzed species.

To account for potential phylogenetic structure, we applied a Brownian Motion (BM) model (Supplementary Fig. 5A), Pagel’s λ regressions — with λ ∈ {0.75 (Supplementary Fig. 5B), 0.50 (Supplementary Fig. 5C), 0.25 (Supplementary Fig. 5D)} — and an Ornstein-Uhlenbeck (OU) selection/adaptation model (Supplementary Fig. 5E). In all five scenarios, the previously observed correlation remained negative and significant (BM: *P =* 0.0057; λ = 0.75: *P =* 0.0029; λ = 0.50: *P =* 0.0024: λ = 0.25: *P =* 0.0021; OU: *P =* 0.0019). In fact, the Akaike information criterion (AIC) indicated that the optimal model was the one that did not incorporate any phylogenetic signal (Supplementary Fig. 5F).

On a more technical note, we also analyzed the consistency of the results using only DMEPs belonging to the aforementioned set of bimodal-shaped probes (12,193 out of 18,268), finding an even stronger negative association (Supplementary Fig. 6A: *R =* −0.7465, *P* = 0.0004, *P_meta_* = 3.43 × 10^−6^), which was also independent of the direction of methylation change (Supplementary Fig. 6B, hyper: 3,666 DMEPs, *R =* −0.7551, *P* = 0.0003, *P_meta_* = 3.28 × 10^−6^; Supplementary Fig. 6C, hypo: 8,247 DMEPs, *R =* −0.5532, *P* = 0.0172, *P_meta_* = 0.0075). Finally, to circumvent differences in normalized age distributions across species, we selected subsets of samples for each species (1,728 samples out of 3,107) so that the distribution of normalized age values was undistinguishable from that of humans (Wilcoxon rank-sum test ≥ 0.05, see Materials and Methods). We then performed the aforementioned correlation analyses using only these subsets, which allowed us to definitively validate the negative associations (DMEPs: *R =* −0.5435, *P* = 0.0197; bimodal DMEPs: *R =* −0.6242, *P* = 0.0056).

Taken together, the results presented in this section indicate that the rate of entropy gain, detectable in the majority of the analyzed CpG sites, is strongly, negatively and linearly associated with mammalian species’ maximum lifespan. This trend remains robust to inaccuracies in reported lifespan values, persists when using time-to-sexual maturity as an alternative life-history trait, is not confounded by phylogenetic relationships, and is validated by analyses restricted to technically reliable probes and to samples with matched normalized age distributions across species.

### Species-specific entropy accumulation over lifetime supports a universal constraint to mammalian longevity

Given the strength of the observed negative association, we aimed to estimate a potential universal upper limit to mammalian lifespan based on the species-specific rate of Shannon entropy accumulation in the DNA methylome with normalized age. To this end, we extrapolated our lineal model to the null entropy rate by determining the cut-off point (X-axis) of the regression line between the entropy gain rate and maximum lifespan (Fig. 5A). This yielded a value of 118 years, which closely aligns with the longest documented lifespan in humans (122.5 years)(1), the species with the highest value for which blood data were available in the Mammalian Methylation Consortium dataset.

**Figure 5.**
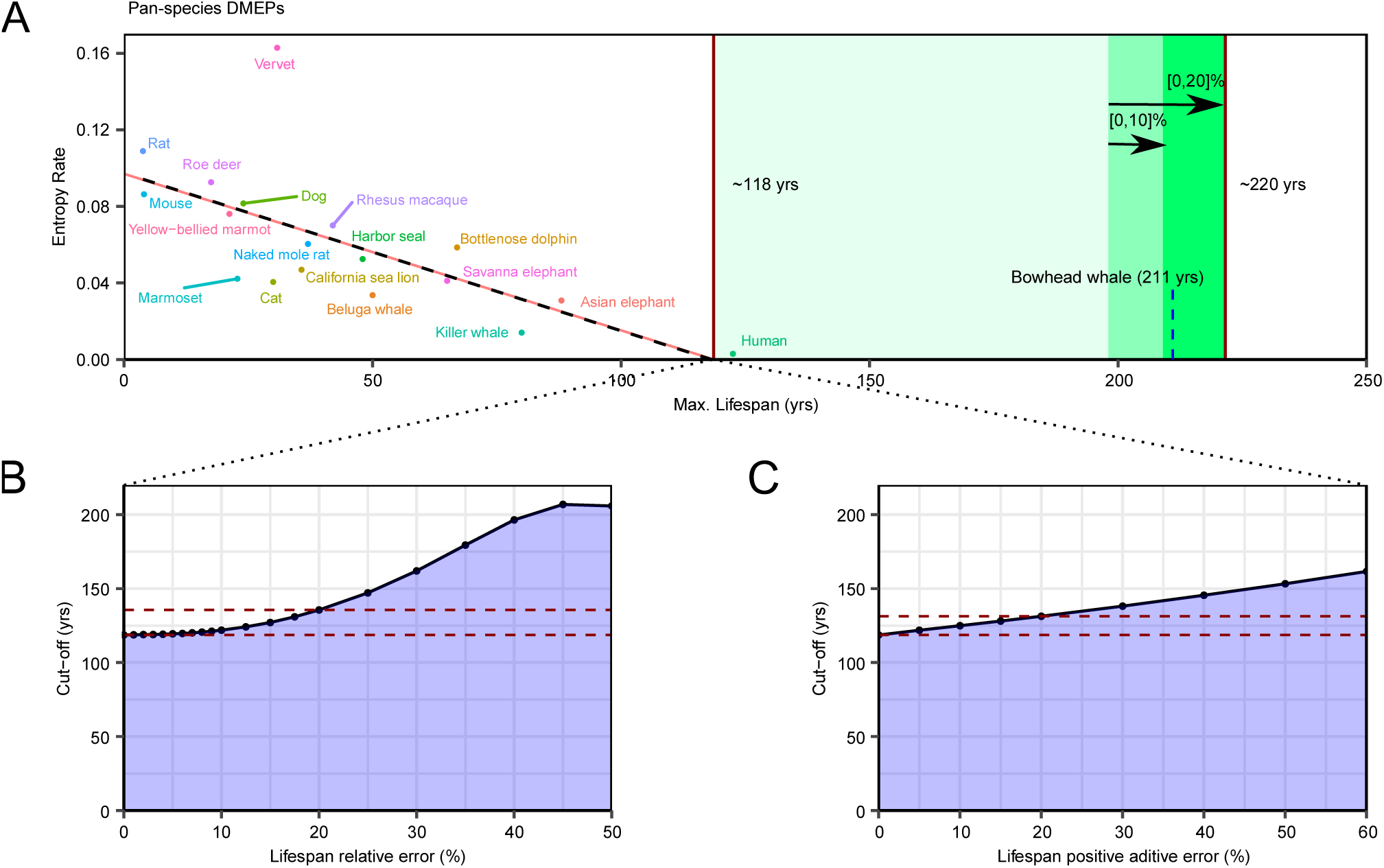
Lifetime rates of epigenetic entropy accumulation reveal a universal mammalian lifespan limit at 220 years. (A) Scatter plot depicting the cut-off point on the X-axis of the regression between entropy gain rate for each species and their respective lifespans, together with the upper bounds derived from different assumptions: either treating the maximum lifespan estimates as fully accurate (198 years), or simulating potential underestimations of up to 10% (209 years) and up to 20% (221 years). The maximum lifespan of the longest-lived mammal, the bowhead whale, is included for comparison. (B-C) Stability dynamics of the X-axis cutoff point under different assumptions: (B) fixed relative errors in lifespan estimates (±*N*%), and (C) varying degrees of lifespan underestimations ([0, +*N*]%).

To assess the robustness of this estimate, we evaluated its sensitivity to potential inaccuracies in reported lifespan values. Introducing fixed relative errors into species’ lifespan estimates tended to shift the X-axis cut-off point upward (Fig. 5B), although changes were negligible up to ±10% (121.91 years) and modest up to ±20% (135.57 years). A similar trend was observed under scenarios of systematic underestimation of lifespan values to varying degrees (Fig. 5C), with the upward shift remaining modest for positive additive errors of + 20%-30% (131.32-138.15 years).

Next, to provide an upper limit for pan-mammalian longevity, we performed empirical simulations (see Materials and Methods) and calculated the 95% confidence interval of the X-axis cut-off point. The resulting range oscillated approximately between 118 and 198 years, under the assumption of perfectly accurate lifespan estimates, and extended to 221 years when accounting for potential underestimations of up to 20% (Materials and Methods). As mentioned earlier, this latter scenario represents a more realistic upper limit for mammalian lifespan, given the uncertainties in species’ estimates, primarily due to differences in sample sizes across studies collecting demographic data and the inherent challenges of obtaining accurate information from predominantly wild species. Notably, this prediction also aligns with current knowledge, as the bowhead whale, the longest-living known mammal, can live up to 211 years(1). Our evidence therefore suggests the existence of a potential pan-mammalian limit to maximum lifespan, which can be inferred from the specific rate of epigenetic noise accumulation over lifetime.

Because aging-associated epigenetic signatures have often been linked to different genomic contexts(25, 26), we explored whether our observations generalized across different chromosomes and gene-context regions. We found that entropy gain rate decreased with mammalian lifespan across all autosomal chromosomes, with the exception of the X chromosome, which did not reach statistical significance (*P =* 0.2133) although it also exhibited a negative linear trend (Supplementary Fig. 7).

Regarding gene-context annotations, regions typically characterized by low DNA methylation levels (promoters, exons and UTRs) exhibited more significant associations in entropy rates with lifespan compared to intergenic and intron regions, which are known to be more susceptible to hypomethylation dysregulation during aging(27) (Supplementary Fig. 8). Taken together, these findings support the generalizability of the observed entropy-lifespan association at the genome-wide level.

### Entropy accumulation over lifetime may explain intersexual differences in mammalian lifespan

Finally, we also investigated whether the epigenetic entropy gain rate could support documented sex differences in lifespan. On average, female lifespan is longer than that of their male counterparts in mammals, but previous studies have found no consistent differences in aging rates(28). To explore this question, we defined epigenetic entropy rate acceleration as the difference in entropy rate between males and females, divided by the rate in females (Supplementary Fig. 9). Out of 17 species tested, 10 showed positive acceleration values, but only 5 species (human, vervet, rat, beluga whale, and Asian elephant) exhibited values of at least +40%. Conversely, of the 7 species with negative acceleration values, only two (mouse and killer whale) were below a -40% difference. Additionally, the same acceleration dynamics across species were observed when using only DMEPs from bimodal shaped probes (Fisher’s exact test, *P =* 5.14 × 10^−5^), and removing sexual-chromosome CpGs did not affect the observed patterns either (Fisher’s exact test, *P =* 5.14 × 10^−5^).

Consequently, it can reasonably be proposed that epigenetic entropy gain rates may provide a potential mechanism to explain the divergence in lifespan between sexes. However, further research is needed to establish conclusive predictors of this phenomenon.

## Discussion

Taken together, our findings have important implications for understanding the range of lifespans observed across mammalian species(1). First, we provide evidence that DNA methylation in evolutionarily conserved genomic regions is maintained more stably with age in long-lived species. Second, we demonstrate that the accumulation of epigenetic noise relative to the lifetime fraction is an informative measure of lifespan differences across mammals. Notably, this association operates independently of the direction of methylation changes and aligns consistently with the information theory of aging(8). It also reflects the process of epigenetic drift(29–31), whereby DNA methylation undergoes subtle, cumulative deviations with age driven by stochastic and environmental factors, leading to individual-specific disordered epigenetic trajectories that appear to progress more rapidly in short-lived species. Third, the discovery of a robust negative linear relationship between noise accumulation and lifespan shifts the focus of future research toward the study of loss of epigenetic information and suggests that epigenetic mechanisms may underlie evolutionary constraints shaping an upper limit of lifespan in mammals. Four, our findings offer molecular evidence that supports recent epidemiological data suggesting human lifespan is nearing a plateau, reinforcing the concept of an intrinsic biological limit to longevity(32).

From a methodological standpoint, we have implemented an independent statistical framework that analyzes the Mammalian array(18) in a global and unbiased manner, using the same CpGs across all samples without prior selections based on probe overlaps or pairwise species comparisons. Nevertheless, our comprehensive approach entails several considerations. First, although methylation data obtained from array-based platforms has provided valuable insights into the study of epigenetic noise across mammalian species, it presents clear limitations when compared to sequencing-based approaches. Specifically, DNA methylation entropy can only be assessed at individual CpG positions, whereas sequencing technologies allow for a more comprehensive analysis of each DNA molecule at two levels: regionally (per read) and at single sites (per CpG, as in arrays)(33, 34). Second, certain age-related changes in DNA methylation, and consequently in epigenetic entropy, are driven by shifts in blood cell-type composition(35–37). The lack of pan-mammalian deconvolution strategies that can be applied across all species studied currently prevents this correction. However, because the observed trends encompass the majority of CpG sites in the Mammalian array and are consistent in different subsets of probes and samples, it is unlikely that this limitation affected the robust conclusions reached in our study. Finally, further research will be needed to elucidate the role of stem cell turnover in determining mammalian lifespan variation, as these divisions are the main factor driving the increase of epigenetic entropy with age in murine tissues(38).

Overall, our results provide compelling evidence that the rate of epigenetic information loss over a lifetime reflects longevity differences across species, and consistently support the existence of a universal lifespan limit in mammalian species of approximately 220 years, first detectable through DNA methylation entropy.

## Materials and Methods

### Data selection and statistical considerations

In line with previous studies(21), we selected blood data from the Mammalian Methylation Consortium(18–20) as the optimal tissue for consistently yielding the highest estimates of lifespan per species. In order to perform pan-species analyses, we also normalized the age of samples using the maximum lifespan of their respective species, thus defining “age/lifespan ratio” or “normalized age”. Subsequently, we excluded blood samples if they (1) belonged to species with fewer than 30 samples, to ensure adequate statistical power to detect robust epigenetic entropy rates; (2) belonged to species with fewer than 5 samples with a normalized age lower than 0.16 (equivalent to 19.60 years in humans); or (3) belonged to species with fewer than 5 samples with a normalized age higher than 0.5 (61.25 years in humans). With these filters, we aimed to select species with a similar normalized age range throughout lifetime and with sufficient representation at all life stages, thus reducing potential biases. After this process, 3,107 blood samples from 18 different species (Fig. 1A) were considered for downstream analyses, covering a wide range of documented mammalian lifespans (from 3.8 to 122.5 years) and several mammalian phylogenetic orders, including 4 primates, 4 rodents, 4 artiodactyls, 4 carnivores and 2 elephants.

### CpG-level differential analyses of entropy and methylation by mammalian species

First, we aimed to identify the evolutionarily conserved CpG sites from Mammalian array (containing 37,488 probes) that gain noise with age in each species. To measure loss of epigenetic information and noise accumulation, we calculated CpG-level Shannon entropy levels from normalized beta values (*β*) according to the following formula(39, 40):

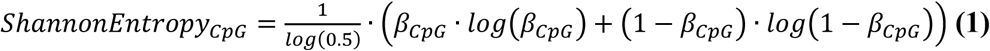

At the CpG level, maximum Shannon entropy (*H =* 1) is achieved when *β =* 0.5, and it decays bidirectionally and non-linearly towards *β =* 0 and *β =* 1. We then employed linear models built with the *limma* package (v.3.44.3)(41) to fit epigenetic entropy values as a function of age in years. Species-specific CpG sites that become noisier with age were defined as those with FDR<0.05 and logFC>0. Furthermore, we performed differential methylation analyses to categorize these CpGs based on their average methylation change, by regressing their corresponding M-values, a logit transformation of beta values that better satisfies the assumptions of linear modeling(42), on age. Accordingly, we classified them as hypermethylated (hyper: FDR<0.05, logFC>0), hypomethylated (hypo: FDR<0.05, logFC<0) or without a clear direction of methylation change (NS: FDR≥0.05).

Subsequently, those CpGs that gain noise with age were linearly correlated with species’ maximum lifespan. To further validate the observed trends and mitigate biases caused by varying numbers of samples per species, which impact statistical power, we computed the residuals by regressing the number of CpG sites that accumulate noise with age against the number of blood samples per species. Finally, we tested the linear associations between residuals and species’ maximum lifespan.

### Pan-species differential analyses of entropy and methylation

In contrast to previous studies that restricted their analyses to gene-associated CpGs shared among species(15) or common sites that change with age in pairwise comparisons(11), we were interested in identifying epigenome-wide longevity signatures through defining pan-mammalian CpGs that gain noise with normalized age. By using the same collection of CpGs across species, we were able to directly compare epigenetic entropy rates without the need to calculate other relative transformations. In addition, this approach enabled the analysis of a larger number of species, as it avoided the need to intersect CpG sites across species, which typically results in a progressively smaller subset.

Consequently, we used the *limma* suite (v.3.44.3) to design multivariate linear mixed models that modeled entropy levels for each CpG as a function of normalized age, while also accounting for sex as a fixed covariate and species as a random-effects component. Our prior selection of 18 species with analogous ranges of normalized age was necessary in order to enable the application of generalized linear models and the derivation of reliable pan-species information. We then defined pan-species differentially methylated entropic positions (DMEPs) as those CpG sites that gain noise with normalized age (FDR<0.05, logFC>0) across mammalian species. Similarly, by performing differential methylation analyses regressing M-values against normalized age (with species and sex as covariates), DMEPs were further classified as hypermethylated (hyper: FDR<0.05, logFC>0), hypomethylated (hypo: FDR<0.05, logFC>0) or without a clear direction of average methylation change (NS: FDR≥0.05).

### Calculations of entropy gain rates and their statistical associations

First, we calculated the accumulation of epigenetic noise occurring in sets of DMEPs for each sample according to the following formula, where *N* is the number of CpG sites:

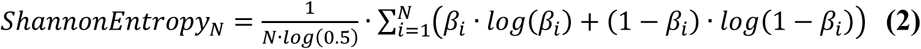

Subsequently, we obtained the epigenetic entropy gain rate per species using the slope of the regression of cumulative entropy levels versus the normalized age of samples. We then investigated the linear correlations between entropy gain rates and two species’ demographic parameters: maximum lifespan and time-to-sexual maturity, revealing a robust negative association in both cases. To account for the uncertainties associated with entropy rate estimates, we applied meta-regression models using the rma function from the *metafor* package (v.4.8.0), incorporating the variance of each rate to validate whether a significant association persists with either species’ maximum lifespan or time-to-sexual maturity.

Finally, we also explored the differences in epigenetic entropy gain rates between sexes by using the slope of the regression of cumulative entropy levels against the normalized age of male or female samples separately. The entropy gain rate acceleration was then defined as the difference between male and female entropy rates divided by the female entropy rate. Acceleration values around ±40% or higher were interpreted as suggestive of meaningful changes.

### Phylogenetic regression modeling

To confirm that our significant associations were not solely driven by phylogenetic inertia, using *nlme* (v3.1.147) and *ape* (v.5.8) packages, we built five distinct phylogenetic regression models that account for the phylogenetic relationships among the 18 species in our study:

(1) A Brownian motion (BM) model, which captures phylogenetic inertia by modeling trait evolution as a random walk with a constant rate of change along the phylogeny.
(2) Three Pagel’s λ models, with λ ∈ {0.75, 0.50, 0.25}, where decreasing values of λ reduce phylogenetic signal strength as expected in Brownian motion model (with λ=1).
(3) An Ornstein-Uhlenbeck (OU) model, which accounts for potential adaptation and selection processes, modeling traits as evolving not only by random fluctuation but also under stabilizing selection toward an optimum.

Finally, we explored which model minimized the Akaike Information Criterion (AIC), including the baseline regression model that did not incorporate any phylogenetic signal.

### Estimation of the universal lifespan limit

To estimate the universal lifespan limit that any mammalian species could potentially reach, we first calculated the cut-off point on the X-axis of the regression line as −*a/b*, where *a* is the intercept and *b* is the slope. We then computed the 95% confidence intervals by simulating 30,000 pairs of slope and intercept values from a multivariate normal distribution, using their original estimates and variance– covariance matrix. Each simulated pair was used to compute a regression line, from which the distribution of X-axis cut-off points was obtained.

Additionally, reported lifespan values exhibit varying levels of uncertainty, primarily due to differences in sample sizes across studies collecting demographic data and the inherent challenges of obtaining accurate information from predominantly wild species, which often leads to underestimation of maximum values. Consequently, we randomly added relative errors to maximum lifespans —ranging from 0% to +10% and from 0% to +20% across species— across 1,000 replicates. Within each replicate, 10,000 slope-intercept pairs were simulated as described above. Finally, 95% confidence intervals were derived from the resulting distributions of X-axis cut-off points.

### Replication of findings using bimodal beta-value probes across mammalian species

Mammalian array was designed for use across multiple mammalian species, but the applicability of its probes decreases as the genome of the species becomes more phylogenetically distant from the human, which is the reference species(43). In fact, density plots of normalized beta methylation values across our 18 selected species revealed a considerable proportion of probes that followed a trimodal-based distribution (Supplementary Fig. 2A), with an intense density peak in β ∈ [0.4,0.6], which may be linked to non-hybridized probes. Because intermediate beta values are associated with maximum DNA methylation information loss, we sought to validate our previous results by eliminating any potential technical bias derived from the array design.

Consequently, we performed a progressive filtering of probes, retaining those that fell outside the range β ∈ [0.4,0.6] in more than a certain percentage of samples (Supplementary Fig. 2B). Stricter filtering (≥ 80% of samples) was sufficient to shift the initial trimodal distribution to a typical bimodal one, with the most stringent threshold (≥95%) producing the best profile. The previous analyses were replicated in full starting only from the selected probes that exhibited a bimodal profile in more than 95% of samples (18,268 CpG sites, 48.73%).

### Replication of findings by modeling uncertainties using bootstrap methods

Entropy rate estimates can be influenced by three main sources of variability: 1) the inclusion of samples within each species, (2) the selection of species used in the correlation of entropy gain rates, and (3) lifespan misestimations. To assess their impact on the robustness of our findings, we first performed a bootstrap analysis with 10,000 replicates by: (a) randomly selecting samples (with replacement) within each species to estimate the slope of the regression between Shannon Entropy and relative age, and (b) randomly selecting species (with replacement) to compute the correlation between entropy gain rates and maximum lifespan. We evaluated the number of significant replicates and the average X-axis cut-off point, revealing that neither sample nor species selection substantially affected the observed significant associations.

To model uncertainties associated with maximum lifespan misestimation, we considered two different approaches. First, within each bootstrap replicate, we applied a gradient of increasing fixed relative errors (from ±1% to ±50%) in lifespan values —randomized in sign for each species— with the purpose of assessing the robustness of findings to directionless perturbations in lifespan estimates. Next, we considered a more realistic scenario in which maximum lifespan was underestimated to distinct degrees. To this end, we randomly added varying relative values to each species’ lifespan estimates —drawn from a uniform distribution between 0% and N_max_(%), where N_max_(%)*=*{5,10,15,20,30,40,50,60}— within each bootstrap replicate. For both approaches, we again evaluated the number of significant replicates and the dynamics of average X-axis cut-off points.

### Replication of findings using normalized age distributions across species

Although species were initially selected to be similar in normalized age ranges, 14 species showed significantly different normalized age distributions compared to humans (the longest living species in the blood methylation dataset), as determined by Wilcoxon-rank sum tests (*P* < 0.05). To address this issue, we designed a selection algorithm that subset samples from each species so that they had a similar normalized age distribution. First, each sample was independently removed from the dataset to calculate differences in mean and standard deviation between the leave-one-out subset of samples and the human reference distribution. Next, samples with normalized age values whose removal minimized mean and standard deviation discrepancies were discarded, and so on until differences were lower than 0.01 for both statistical parameters. After applying this algorithm to the 14 species, the selected subsets of samples exhibited undistinguishable normalized age distributions (*P ≥* 0.05) from humans. We again replicated previous analyses using these new subsets of samples.

### Visualization of results

All graphs were generated in R using the *ggplot2* package (v.3.3.2)(44).

## Acknowledgments

This work was supported by the Spanish Association Against Cancer (PROYE18061FERN and PRYGN235109FERN to MFF), the Asturias Government (PCTI) co-funding 2018-2023/FEDER (IDI/2018/146 and IDI/2021/000077 to MFF), the ISCIII (PI21/01067 to MFF and AFF, COV00624 to JRT and MFF), CIBERER Acciones Cooperativas y Complementarias Intramurales (ACCI20-34-U766 to MFF), ISPA and the Galbán Association (2023-165-GALBAN-TEVAJ to JRT), ISPA-Jannsen (2021-048-INTRAMURAL NOV-TEVAR to JRT), CSIC (202020E092 to MFF), and the European Commission NextGenerationEU, through CSIC’s Global Health Platform (PTI Salud Global) and the Spanish Ministry of Science and Innovation through the Recovery, Transformation and Resilience Plan (SGL2021-03-39 and SGL2021-03-040). JJAL is supported by the Spanish Association Against Cancer (Grant number PRDAS21642ALBA), and JRT by a Ramon y Cajal fellowship from the Spanish Ministry of Science and Innovation (RYC2021-031799-I). We also acknowledge support from the Institute of Oncology of Asturias (IUOPA, supported by Obra Social Cajastur-Liberbank, Spain), the Health Research Institute of Asturias (ISPA-FINBA) and the Consorcio Centro de Investigación Biomédica en Red de Enfermedades Raras (CIBERER-ISCIII).

## Author contributions

JJAL, RFP and MFF conceived, coordinated and supervised the study. JJAL designed and performed the main data analyses. JRT and AFF assisted with the interpretation of results. JJAL, RFP and MFF participated in drafting the manuscript. All authors revised, read and approved the final manuscript.

## Supplementary Figures

**Supplementary Fig. 1.**
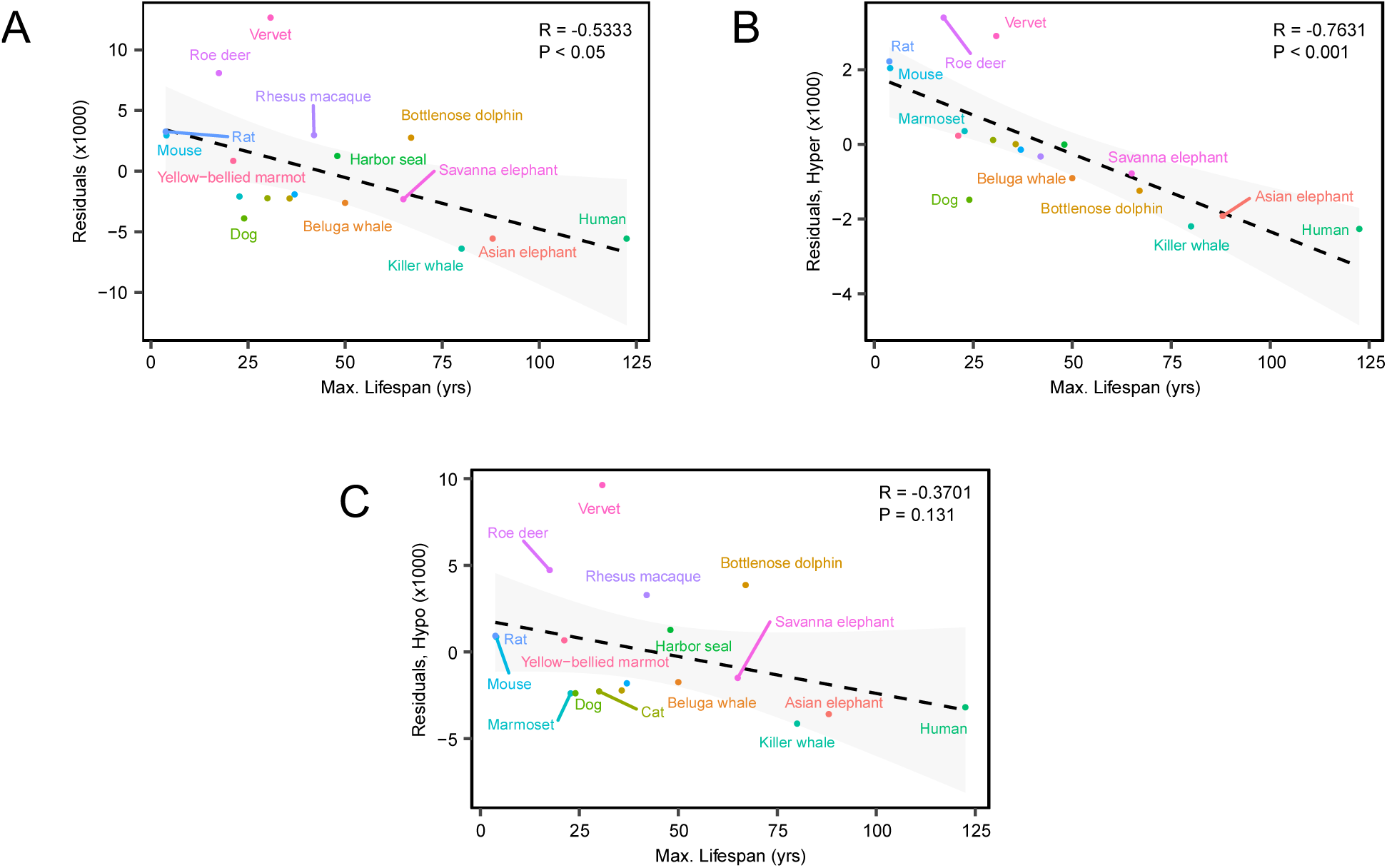
Scatter plots depicting the relationship between species’ maximum lifespan and residuals of the linear regression between the number of CpG sites that accumulate noise with age and the number of samples per species (A), further separating by hyper- (B) and hypomethylation (C) mechanisms.

**Supplementary Fig. 2.**
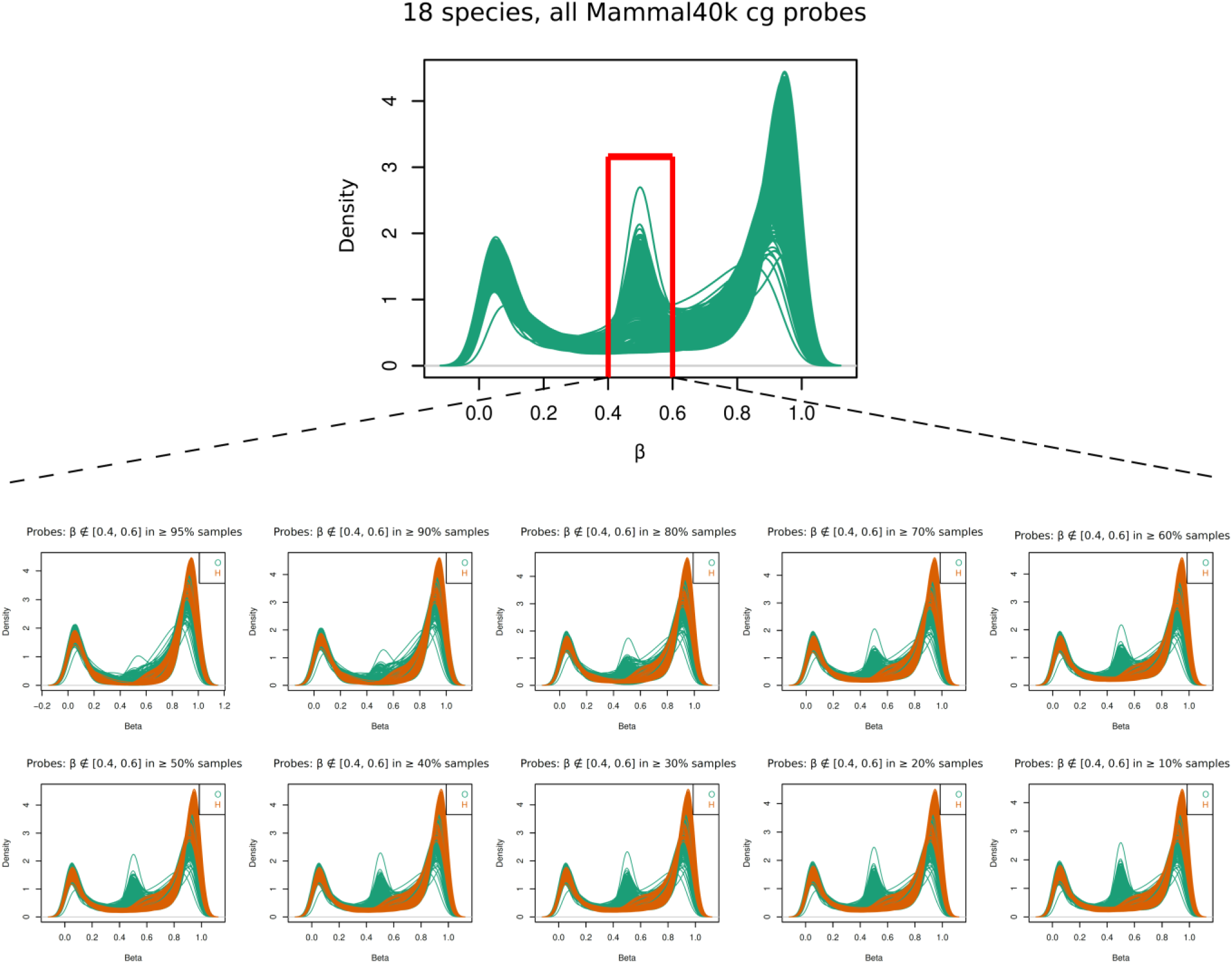
Up, Density plots showing the distribution of DNA methylation beta values across the Mammalian array, with the peak of potentially undetected/problematic probes at [0.4, 0.6] (highlighted in red). Below, probe filtering of these probes based on different sample thresholds has been applied to obtain density profiles with a bimodal shape (human reference samples are colored in orange).

**Supplementary Fig. 3.**
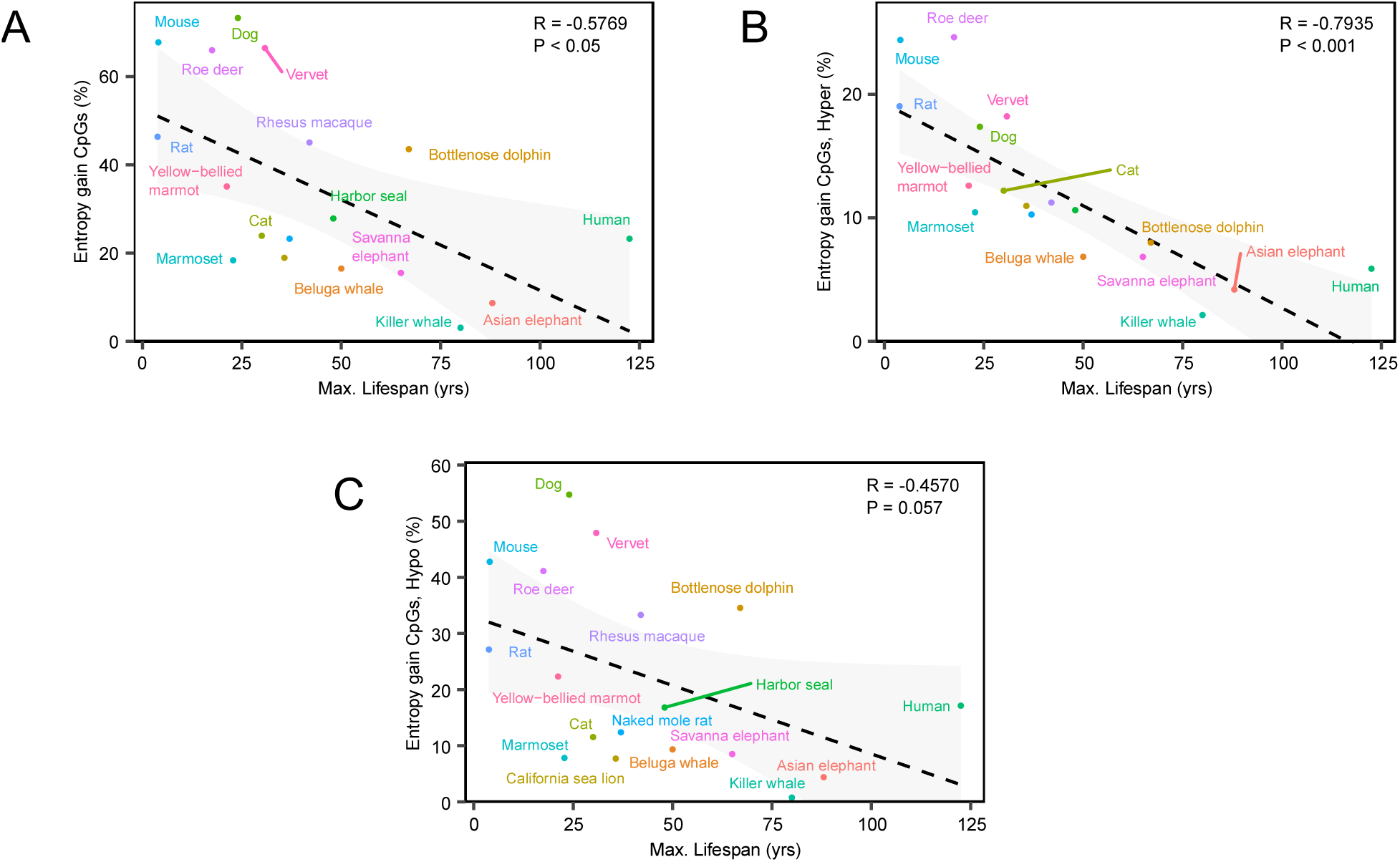
Scatter plots showing the correlation between the number of bimodal-shaped CpG sites that accumulate noise with age and species’ maximum lifespan (A), further separating by hyper- (B) and hypomethylation (C) processes.

**Supplementary Fig. 4.**
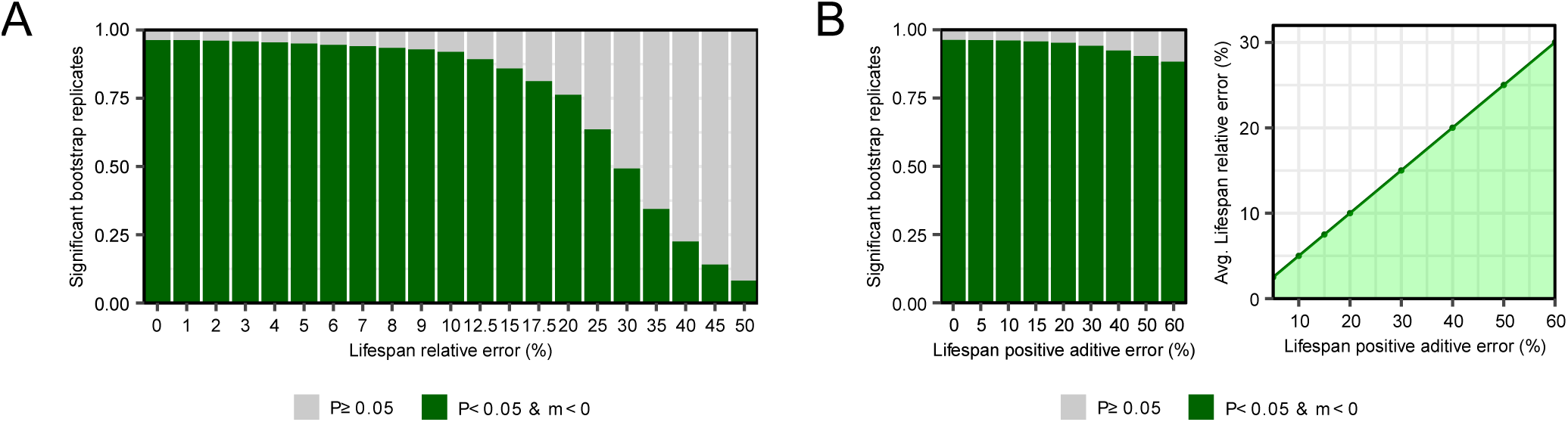
Barplots showing the proportion of bootstrap replicates with a negative and significant association between entropy gain rates and species’ maximum lifespan, under assumptions of (A) fixed relative errors in lifespan estimates (±*N*%), or (B) varying degrees of lifespan underestimations ([0, +*N*]%). Bars at 0% represent the effect of sample and species resampling without introducing perturbations in lifespan estimates. Supplementary Fig. 4B also includes a correlation to compare both lifespan misestimation scenarios.

**Supplementary Fig. 5.**
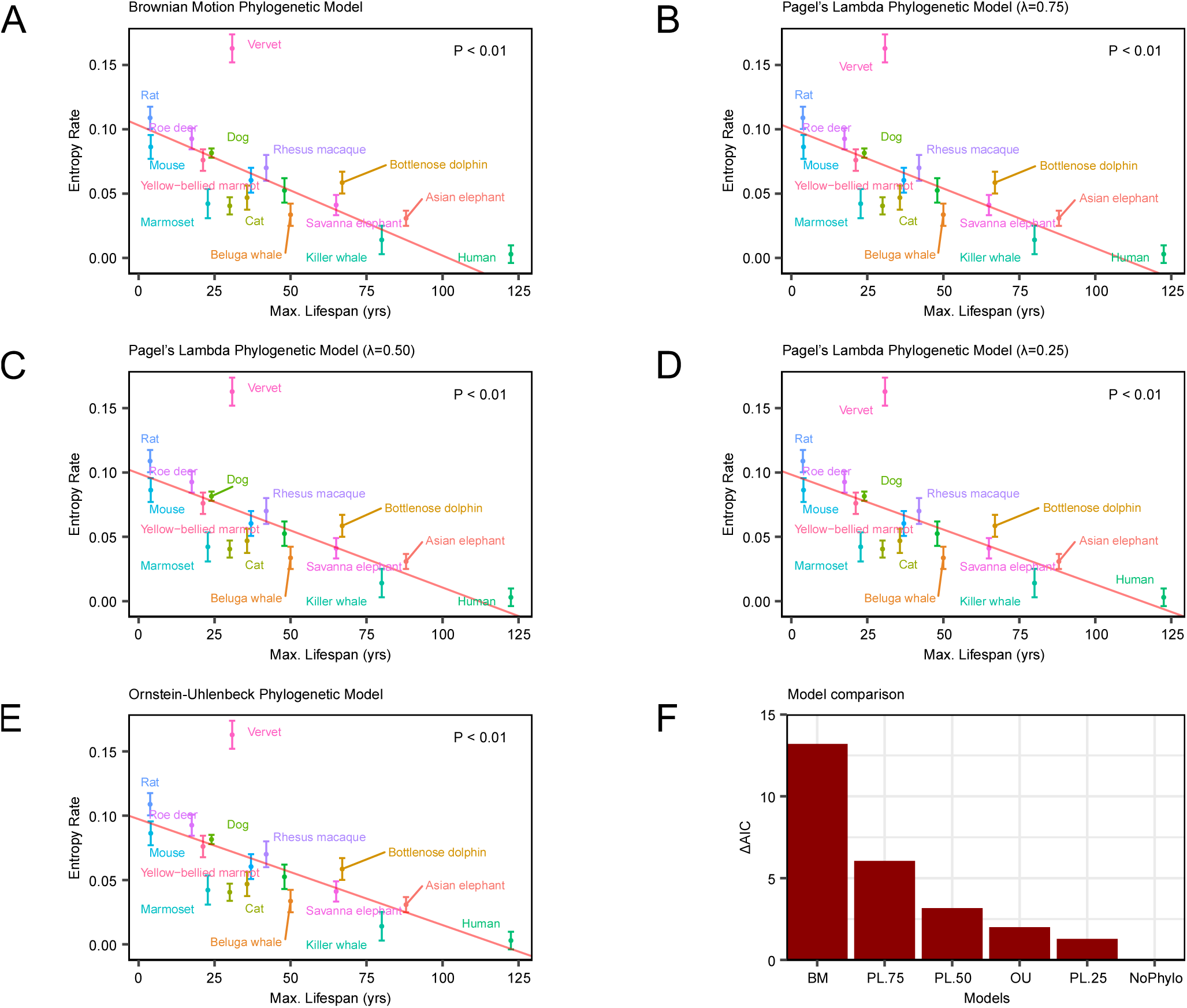
Scatter plots showing distinct phylogenetic regression models (in red) between entropy gain rates and species’ maximum lifespans based on: (A) a Brownian motion model; (B-D) three Pagel’s lambda models moderating phylogenetic signal, and (E) a Ornstein-Uhlenbeck selection/adaptation model. Standard errors for each rate are represented with bars at each point. (F) Barplots showing the delta Akaike Information Criterion (ΔAIC) values for all tested scenarios, including a non-phylogenetic model, to determine the best-fitting via AIC minimization.

**Supplementary Fig. 6.**
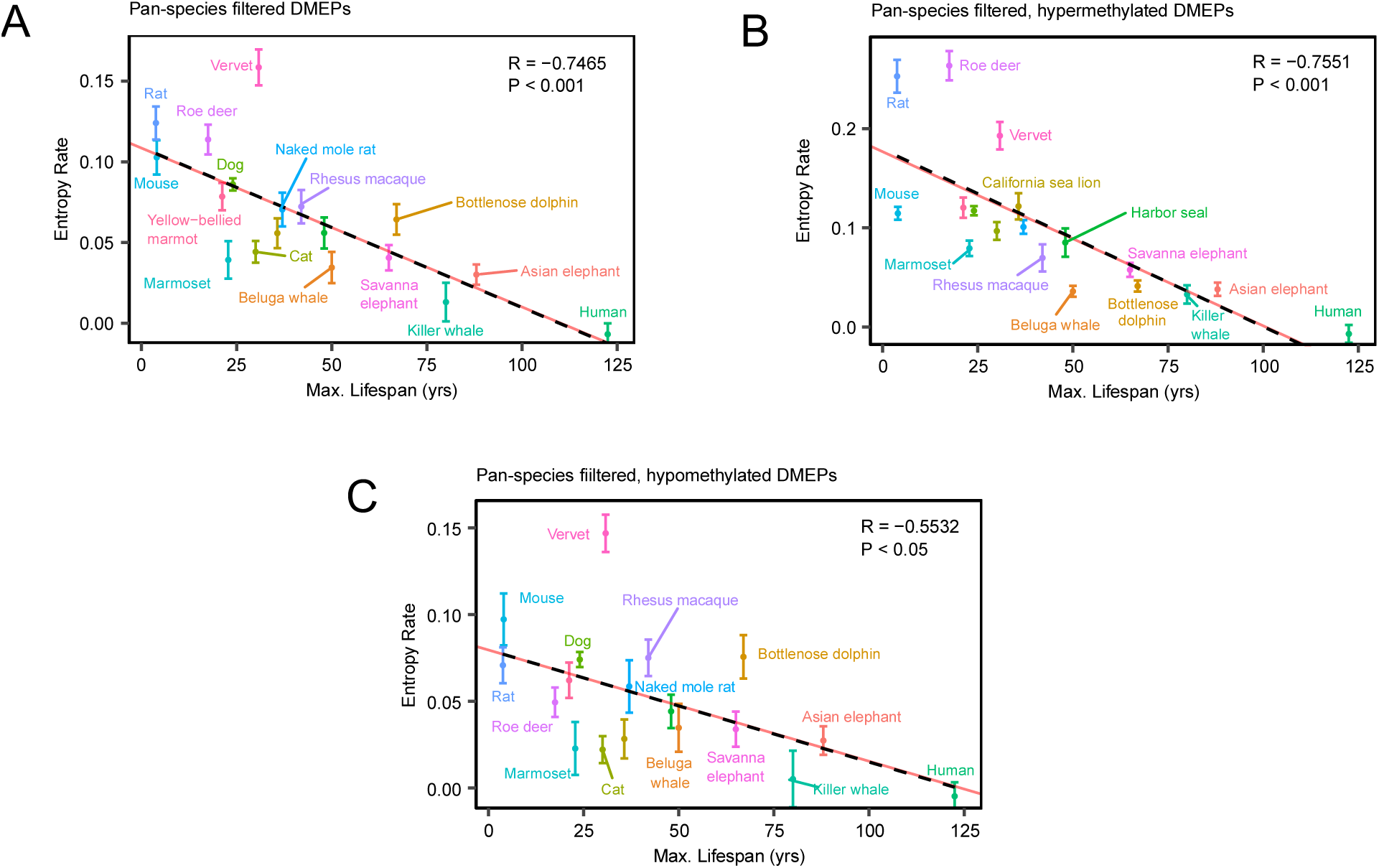
Scatter plots depicting the linear decrease in epigenetic entropy gain rate with age in bimodal-shaped probes as a function of species’ maximum lifespan (A), further separating into hyper- (B) and hypomethylation (C) processes. Standard errors for each rate are included, along with meta-analysis regression lines (shown in red).

**Supplementary Fig. 7.**
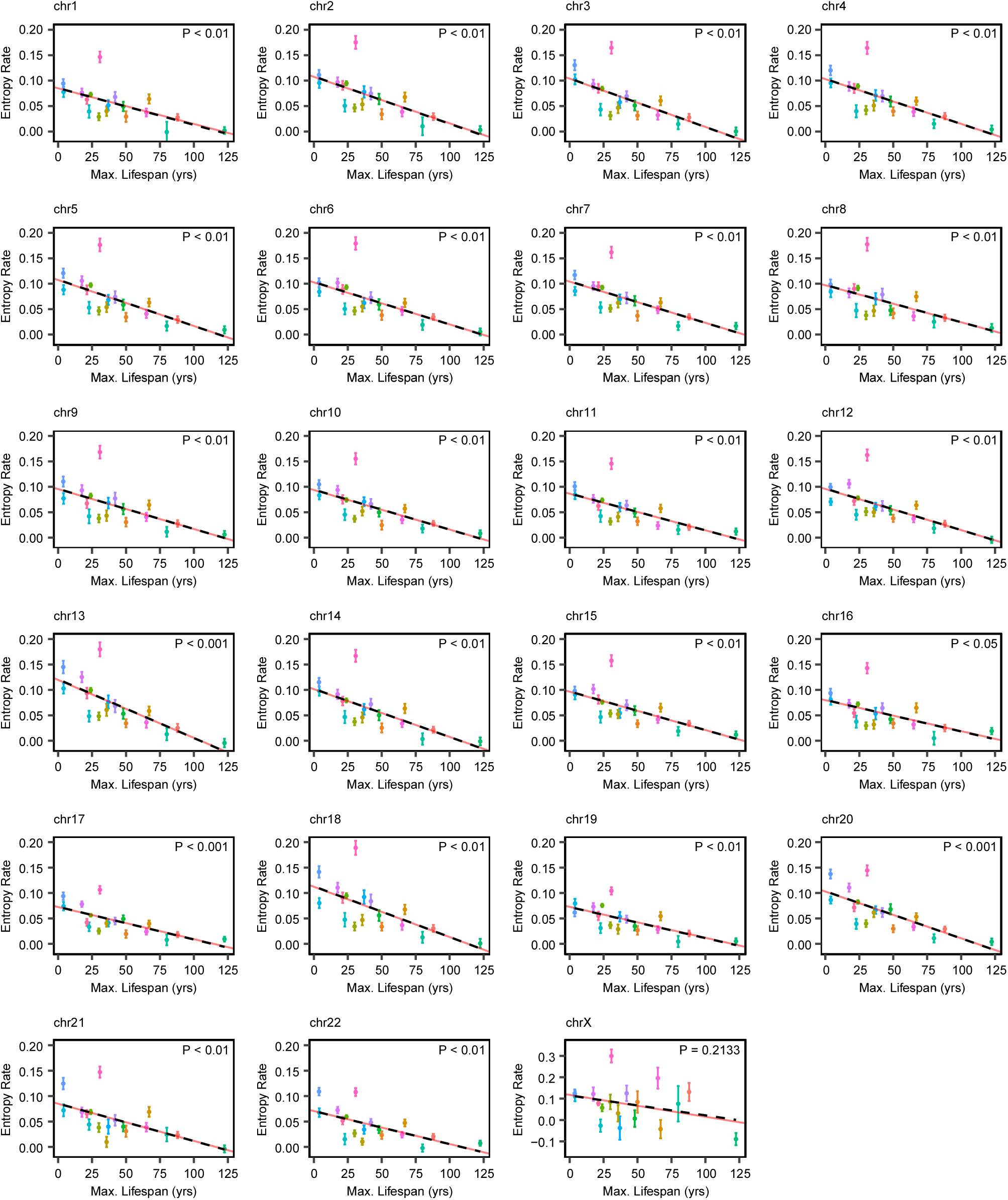
Scatter plots representing the relationship between epigenetic entropy gain rate and species’ maximum lifespan across different human chromosomes. Standard errors for each rate are shown along with meta-analysis regression lines (in red).

**Supplementary Fig. 8.**
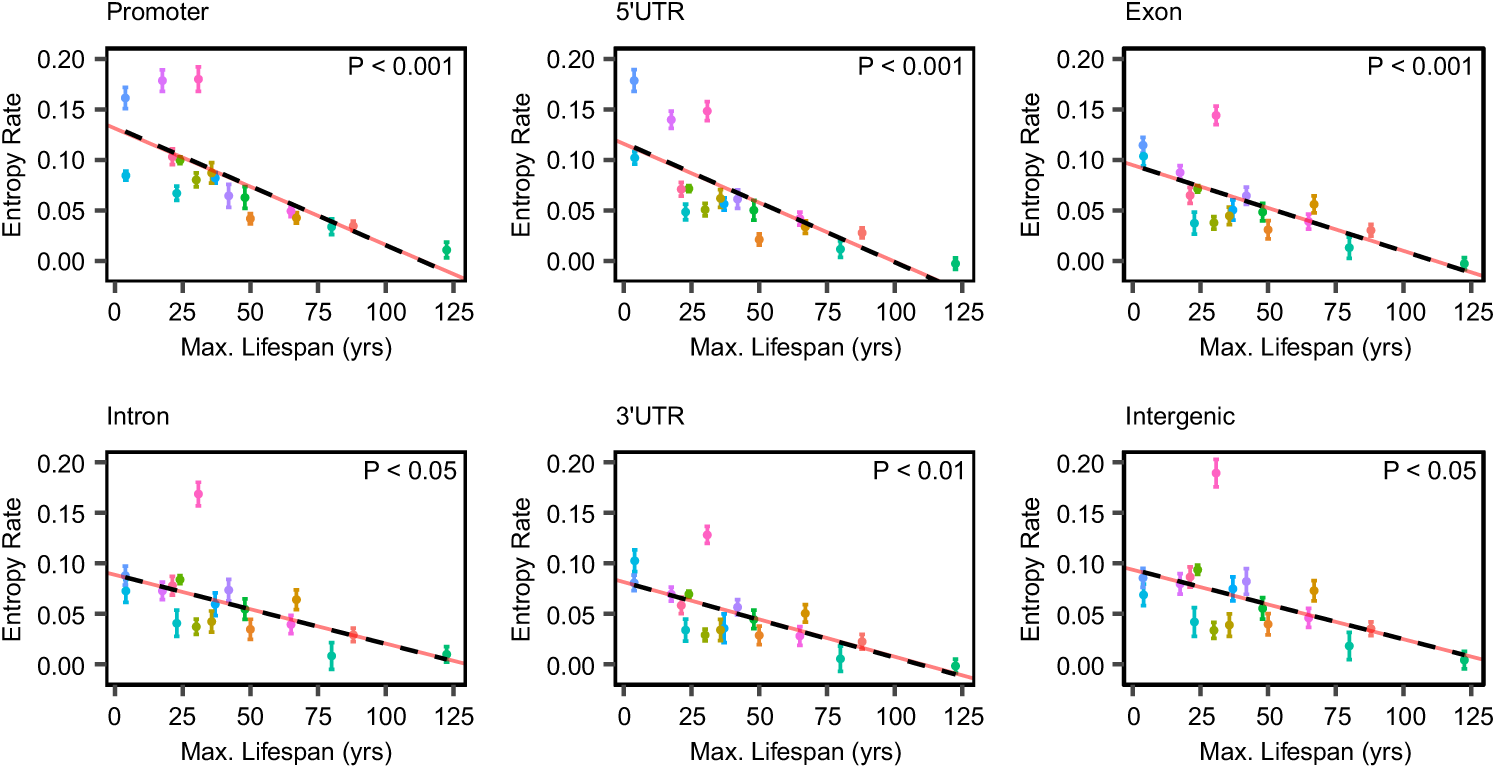
Scatter plots depicting the association between epigenetic entropy gain rate and species’ maximum lifespan across different gene-context regions. Standard errors for each rate are included, along with meta-analysis regression lines (in red).

**Supplementary Fig. 9.**
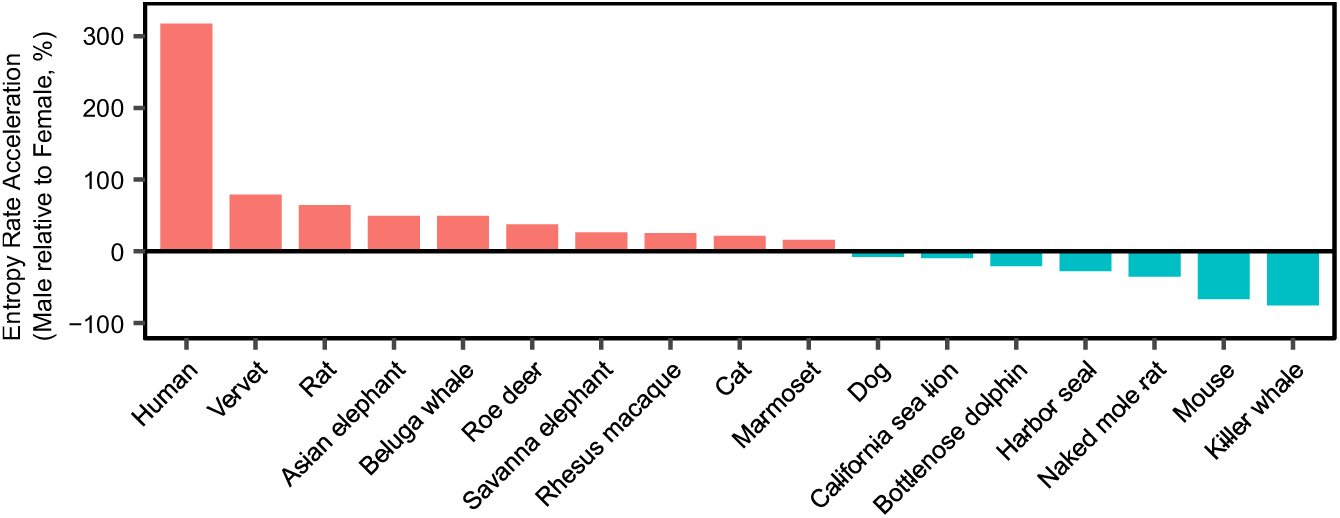
Barplots showing the dynamics of epigenetic entropy rate acceleration (males relative to females) across different mammalian species.

## References

1. J. P. De Magalhães, J. Costa, A database of vertebrate longevity records and their relation to other life-history traits. Journal of Evolutionary Biology 22, 1770–1774 (2009).

2. M. T. Mc Auley, The evolution of ageing: classic theories and emerging ideas. Biogerontology 26, 6 (2024).

3. V. N. Gladyshev, Aging: progressive decline in fitness due to the rising deleteriome adjusted by genetic, environmental, and stochastic processes. Aging Cell 15, 594–602 (2016).

4. X. Tian, A. Seluanov, V. Gorbunova, Molecular Mechanisms Determining Lifespan in Short- and Long-Lived Species. Trends in Endocrinology & Metabolism 28, 722–734 (2017).

5. J.-Y. Lee, et al., Misexpression of genes lacking CpG islands drives degenerative changes during aging. Science Advances 7, eabj9111 (2021).

6. H. Izgi, et al., Inter-tissue convergence of gene expression during ageing suggests age-related loss of tissue and cellular identity. eLife 11, e68048 (2022).

7. P. Sen, et al., Spurious intragenic transcription is a feature of mammalian cellular senescence and tissue aging. Nat Aging 3, 402–417 (2023).

8. Y. R. Lu, X. Tian, D. A. Sinclair, The Information Theory of Aging. Nat Aging 3, 1486–1499 (2023).

9. J.-H. Yang, et al., Loss of epigenetic information as a cause of mammalian aging. Cell 186, 305–326.e27 (2023).

10. A. Cagan, et al., Somatic mutation rates scale with lifespan across mammals. Nature 604, 517–524 (2022).

11. S. J. C. Crofts, E. Latorre-Crespo, T. Chandra, DNA methylation rates scale with maximum lifespan across mammals. Nat Aging 4, 27–32 (2024).

12. S. Horvath, J. Zhang, A. Haghani, A. T. Lu, Z. Fei, Fundamental equations linking methylation dynamics to maximum lifespan in mammals. Nat Commun 15, 8093 (2024).

13. V. K. Rakyan, et al., Human aging-associated DNA hypermethylation occurs preferentially at bivalent chromatin domains. Genome Res. 20, 434–439 (2010).

14. M. A. Koldobskiy, O. Camacho, P. Reddy, J. C. I. Belmonte, A. P. Feinberg, Convergence of aging- and rejuvenation-related epigenetic alterations on PRC2 targets. [Preprint] (2023). Available at: https://www.biorxiv.org/content/10.1101/2023.06.08.544045v1 [Accessed 8 July 2025].

15. E. M. Bertucci-Richter, B. B. Parrott, The rate of epigenetic drift scales with maximum lifespan across mammals. Nat Commun 14, 7731 (2023).

16. K. Seale, S. Horvath, A. Teschendorff, N. Eynon, S. Voisin, Making sense of the ageing methylome. Nat Rev Genet 23, 585–605 (2022).

17. Y. Fang, et al., DNA methylation entropy is associated with DNA sequence features and developmental epigenetic divergence. Nucleic Acids Research 51, 2046–2065 (2023).

18. A. Arneson, et al., A mammalian methylation array for profiling methylation levels at conserved sequences. Nat Commun 13, 783 (2022).

19. A. Haghani, et al., DNA methylation networks underlying mammalian traits. Science 381, eabq5693 (2023).

20. A. T. Lu, et al., Universal DNA methylation age across mammalian tissues. Nat Aging 3, 1144–1166 (2023).

21. C. Z. Li, et al., Epigenetic predictors of species maximum life span and other life-history traits in mammals. Science Advances 10, eadm7273 (2024).

22. S. M. Baylis, M. de Lisle, M. E. Hauber, Inferring maximum lifespan from maximum recorded longevity in the wild carries substantial risk of estimation bias. Ecography 37, 770–780 (2014).

23. J. P. de Magalhães, J. Costa, G. M. Church, An Analysis of the Relationship Between Metabolism, Developmental Schedules, and Longevity Using Phylogenetic Independent Contrasts. The Journals of Gerontology: Series A 62, 149– 160 (2007).

24. R. E. Ricklefs, Life-history connections to rates of aging in terrestrial vertebrates. Proceedings of the National Academy of Sciences 107, 10314–10319 (2010).

25. C. Huidobro, A. F. Fernandez, M. F. Fraga, Aging epigenetics: Causes and consequences. Molecular Aspects of Medicine 34, 765–781 (2013).

26. S. Pal, J. K. Tyler, Epigenetics and aging. Science Advances 2, e1600584 (2016).

27. R. F. Pérez, J. R. Tejedor, G. F. Bayón, A. F. Fernández, M. F. Fraga, Distinct chromatin signatures of DNA hypomethylation in aging and cancer. Aging Cell 17, e12744 (2018).

28. J.-F. Lemaître, et al., Sex differences in adult lifespan and aging rates of mortality across wild mammals. Proceedings of the National Academy of Sciences 117, 8546–8553 (2020).

29. M. F. Fraga, et al., Epigenetic differences arise during the lifetime of monozygotic twins. Proceedings of the National Academy of Sciences 102, 10604–10609 (2005).

30. Q. Tan, et al., Epigenetic drift in the aging genome: a ten-year follow-up in an elderly twin cohort. International Journal of Epidemiology 45, 1146–1158 (2016).

31. C. A. Reynolds, et al., A decade of epigenetic change in aging twins: Genetic and environmental contributions to longitudinal DNA methylation. Aging Cell 19, e13197 (2020).

32. J. Andrade, C. G. Camarda, H. Pifarré i Arolas, Cohort mortality forecasts indicate signs of deceleration in life expectancy gains. Proceedings of the National Academy of Sciences 122, e2519179122 (2025).

33. E. M. Bertucci-Richter, E. P. Shealy, B. B. Parrott, Epigenetic drift underlies epigenetic clock signals, but displays distinct responses to lifespan interventions, development, and cellular dedifferentiation. Aging 16, 1002–1020 (2024).

34. D. A. Landau, et al., Locally Disordered Methylation Forms the Basis of Intratumor Methylome Variation in Chronic Lymphocytic Leukemia. Cancer Cell 26, 813–825 (2014).

35. Z. Zhang, et al., Deciphering the role of immune cell composition in epigenetic age acceleration: Insights from cell-type deconvolution applied to human blood epigenetic clocks. Aging Cell 23, e14071 (2023).

36. H. Tong, et al., Cell-type specific epigenetic clocks to quantify biological age at cell-type resolution. Aging 16, 13452– 13504 (2024).

37. K. Seale, A. Teschendorff, A. P. Reiner, S. Voisin, N. Eynon, A comprehensive map of the aging blood methylome in humans. Genome Biology 25, 240 (2024).

38. H. Vaidya, et al., DNA methylation entropy as a measure of stem cell replication and aging. Genome Biology 24, 27 (2023).

39. G. Hannum, et al., Genome-wide Methylation Profiles Reveal Quantitative Views of Human Aging Rates. Molecular Cell 49, 359–367 (2013).

40. C. E. Shannon, W. Weaver, Mathematical Theory of Communication (1963).

41. M. E. Ritchie, et al., limma powers differential expression analyses for RNA-sequencing and microarray studies. Nucleic Acids Res 43, e47 (2015).

42. P. Du, et al., Comparison of Beta-value and M-value methods for quantifying methylation levels by microarray analysis. BMC Bioinformatics 11, 587 (2010).

43. W. Ding, D. Kaur, S. Horvath, W. Zhou, Comparative epigenome analysis using Infinium DNA methylation BeadChips. Briefings in Bioinformatics 24, bbac617 (2023).

44. H. Wickham, ggplot2: Elegant Graphics for Data Analysis (Springer, 2009).

